# A dense linkage map of Lake Victoria cichlids improved the *Pundamilia* genome assembly and revealed a major QTL for sex-determination

**DOI:** 10.1101/275388

**Authors:** Philine G.D. Feulner, Julia Schwarzer, Marcel P. Haesler, Joana I. Meier, Ole Seehausen

## Abstract

Genetic linkage maps are essential for comparative genomics, high quality genome sequence assembly and fine scale quantitative trait locus (QTL) mapping. In the present study we identified and genotyped markers via restriction-site associated DNA (RAD) sequencing and constructed a genetic linkage map based on 1,597 SNP markers of an interspecific F2 cross of two closely related Lake Victoria cichlids (*Pundamilia pundamilia* and *P.* sp. “red head”). The SNP markers were distributed on 22 linkage groups and the total map size was 1,594 cM with an average marker distance of 1.01 cM. This high-resolution genetic linkage map was used to anchor the scaffolds of the *Pundamilia* genome and estimate recombination rates along the genome. Via QTL mapping we identified a major QTL for sex in a ∼1.9 Mb region on Pun-LG10, which is homologous to *Oreochromis niloticus* LG 23 (Ore-LG23) and includes a well-known vertebrate sex-determination gene (*amh*).

## Introduction

The haplochromine cichlid lineage of the East African Great Lakes is famous for forming large adaptive radiations often in exceptionally short time, resulting in several hundred species each in Lakes Malawi and Victoria, and dozens of species each in several smaller East African Lakes (Kocher 2004; Seehausen 2015). The Lake Victoria haplochromine cichlid radiation stands out in being the youngest (∼15,000 years) showing a high degree of diversity in morphology, behavior and ecology (Greenwood 1974; Seehausen 1996). An abundance of studies have been published on the evolution of Lake Victoria cichlids, providing insight to colonization history (Nagl *et al.* 2000; Seehausen *et al.* 2003; Verheyen *et al.* 2003; Meier *et al.* 2017b), species formation (Seehausen *et al.* 1997; Seehausen and van Alphen 1999; Seehausen *et al*. 1999; Selz *et al.* 2014a), the interaction of sexual and natural selection (Seehausen 2000, Seehausen *et al.* 2008, Maan and Seehausen 2011), and the role of hybridization between distant relatives (Seehausen *et al.* 2003, Keller *et al.* 2013, Selz *et al.* 2014b, Meier *et al.* 2017a, Meier *et al.* 2017b). Recently, several cichlid genomes were published (Brawand *et al*. 2014), among them one from Lake Victoria. This genome has been started to be used to investigate the genomic landscape of speciation (Meier *et al.* accepted). Detailed genetic linkage maps offer a powerful tool to improve the quality of genome assemblies (Fierst 2015) and set the framework for quantitative trait loci (QTL) localization. In the past decade, a number of genetic linkage maps have been published for haplochromine cichlids using various molecular genetic markers (Streelman *et al.* 2003; Sanetra *et al.* 2009; O´Quin *et al.* 2013; Henning *et al.* 2014; 2017). For Lake Victoria cichlids three linkage maps based on two interspecific F2 hybrid crosses were published. The first was based on an F2 cross between *Paralabidochromis chilotes* and *Paralabidochromis sauvagei* and contained 184 microsatellites and two SNP markers with a mean marker spacing of 6.09 cM on 25 linkage groups (Kudo *et al.* 2015). The two others were based on F2 crosses between *Paralabidochromis sauvagei* and *Pundamilia* cf. *nyererei* (Henning *et al*. 2014) and *Paralabidochromis chilotes* and *Pundamilia* cf. *nyererei* (Henning *et al*. 2017). Linkage maps were constructed with 867 and 752 single-nucleotide polymorphism (SNP) markers resulting in a mean marker spacing of 1.30 and 1.09 cM, respectively on 22 linkage groups (Henning *et al.* 2014; 2017). These linkage maps were then used to identify QTL, such as for lateral stripes, lip size, and head morphology (Henning *et al*. 2014; 2017) and sex determination (Kudo *et al*. 2015). None of the linkage maps has been used to improve the Lake Victoria haplochromine genome assembly.

In haplochromine cichlids, some polymorphic color patterns are genetically linked to sex determination and are associated with segregating polymorphisms in sex determination (Holzberg 1978; Seehausen *et al*. 1999; Lande *et al*. 2001; Streelman *et al*. 2003; Kocher 2004). These observations supported the hypothesis that the rapid evolution of sex determination systems might play a role in the very rapid speciation of haplochromine cichlids (Seehausen *et al*. 1999; Lande *et al*. 2001; Kocher 2004; Ser *et al*. 2010). A high diversity of sex determination systems and high sex chromosome turnover rates are known in fish, including cichlids, with a variety of environmental and genomic factors resulting in male or female phenotypes (reviewed e.g. in Heule *et al*. 2014a). In cichlids, very closely related species, populations within the same species, and even individuals within a population, can have different sex determination mechanisms or non-homologous sex chromosomes. This is evidenced by the presence of both XX-XY and ZZ-ZW sex determination systems within haplochromines of Lakes Victoria and Malawi and in oreochromine cichlids (Seehausen *et al*. 1999; Lande *et al*. 2001; Cnaani *et al*. 2008; Roberts *et al*. 2009; Ser *et al*. 2010). Some candidates for genetic sex determination in cichlids exist and could be associated with respective chromosomes. Among different species of *Oreochromis*, sex determination loci have been repeatedly mapped on linkage group (LG) 1 (XY), LG 3 (ZW) and LG 23 (XY) (Cnaani *et al*. 2008) and in haplochromine cichlids, sex determination loci mainly mapped to LG 5 (ZW and XY) and LG 7 (XY) (Ser *et al*. 2010; Kudo *et al*. 2015; Roberts *et al*. 2016; Böhne *et al*. 2016; Peterson *et al*. 2017). Some genes that have repeatedly evolved as master sex determination genes in teleost fishes (Kikuchi and Hamaguchi 2013; Heule *et al*. 2014a) seem to play a role in sex determination in cichlids as well. Recent results published on *Astatotilapia calliptera*, a haplochromine cichlid from Lake Malawi, and *Oreochromis niloticus*, a distant relative of the East African adaptive radiations, indicate that two of these candidate genes, the gonadal soma-derived factor (*gsdf*) and the anti Müllerian hormone (*amh*) might have been re-used as sex determination loci (Eshel *et al*. 2014; Peterson *et al*. 2017). Those genes are often derived by duplication or allelic diversification from genes with a known function in sex differentiation or gonad development (Heule *et al*. 2014a).

In the present study we construct a linkage map of an interspecific F2 cross between two very closely related Lake Victoria cichlid species (*Pundamilia pundamilia* and *P.* sp. “red head). The map was build using 1,597 SNPs identified and genotyped via restriction-site associated DNA (RAD) sequencing with an average marker distance of 1.01 cM. We then used the linkage map to anchor the scaffolds of the *P. nyererei* reference genome to the 22 linkage groups of the map and to perform a QTL analysis for putative sex determination loci in *Pundamilia*. We identify the LG determining sex in a Lake Victoria cichlid cross, as well as potential candidate genes for sex determination and put these findings into the context of sex determination evolution within a rapidly radiating clade of fish.

## Materials and Methods

### Mapping family and RAD sequencing

The genetic cross was started with a lab bred *Pundamilia* sp. “red head” male from Zue Island in Lake Victoria (lab strain established from wild caught fishes by OS in 1993, 4^th^ or 5th lab generation) and a wild *P. pundamilia* female caught by OS at Makobe Island in Lake Victoria in 2003. Eggs were removed from the female’s mouth five days after spawning and reared in isolation from the adults. After reaching maturity, four F1 individuals were crossed, resulting in two F2 families with together more than 300 individuals. When F2 individuals were adult and sexually mature sex was determined based on coloration, then fish were euthanized with MS222, and a fin clip was taken and stored in 98% ethanol for genetic analyses. Genomic DNA of 218 F2 progeny, the four F1 parents, and the two F0 grandparents was extracted using phenol-chloroform. Restriction-site associated DNA (RAD) sequencing libraries were prepared following Marques *et al*. (2016) using a protocol slightly modified from Baird *et al*. (2008). In brief, genomic DNA was digested with *Sbf*I followed by shearing and size selection of 300 to 500 bp. Equimolar proportions of DNA from 11 to 48 individuals carrying different barcode sequences were pooled into one library. Each library was amplified in four reactions of 50 µl aliquots. A total of nine libraries were single-end sequenced (100 bp) each on a single lane of an Illumina HighSeq 2500 platform either at the Next Generation Sequencing Platform of the University of Bern or at the Genomic Technologies Facility of the University of Lausanne. Some individuals and all F0 grandparents were sequenced in up to three libraries to increase coverage. Together with each library, we sequenced about 10% reads of bacteriophage PhiX genomic DNA (Illumina Inc.) to increase complexity at the first 10 sequenced base pairs. During read processing, PhiX reads were further utilized to recalibrate libraries to equalize base quality scores across Illumina lanes utilizing GATK version 3.2 (McKenna *et al*. 2010).

### Sequence processing and genotyping

Before recalibration, read qualities were inspected using fastQC (http://www.bioinformatics.babraham.ac.uk/projects/fastqc) and filtered using FASTX Toolkit 0.0.13 (http://hannonlab.cshl.edu/fastx_toolkit/index.html) requiring a minimum quality of 10 at all bases and of 30 in at least 95% of the read. After PhiX removal, reads were demultiplexed, cleaned, and trimmed to 92 bp with process_radtags implemented in Stacks v1.26 (Catchen *et al*. 2013). Reads were mapped against the *P. nyererei* reference genome (Brawand *et al*. 2014) using bowtie2 version 2.2.6 (Langmead and Salzberg 2012). Mapped reads of individuals run in multiple libraries were merged using Picard tools version 1.97 and filtered for a mapping quality of at least 30. After the filtering pipeline we were left with a total of 719,720,265 sequences across the nine RAD libraries (on average 79,970,000 reads per library). For the female and male parental samples, 1.364,225 and 6,459,242 reads respectively, were mapped and remained after filtering. For the 222 progeny individuals (including the F1) we obtained on average 2,008,826 reads per individual. All 224 individuals (218 F2, the two grandparents and four F1) were genotyped using freebayes version 1.0.0 (Garrison and Marth 2012). As a first filter, sites were kept if bi-allelic, had less then 50% missing data, a quality of more than 2, a minor allele frequency of more than 5%, and a minimal depth of 3. Utilising a script established to filter freebayes genotype calls based on RAD sequencing (https://github.com/jpuritz/dDocent/blob/master/scripts/dDocent_filters), genotypes were further excluded (thresholds given in brackets) on criteria related to allelic balance at heterozygote sites (< 0.28 allele balance between reads), quality versus depth (ratio <0.5), strand presentation (overlapping forward and reverse reads), and site depth (one standard deviation from mean and a quality score lower than twice the depth first, followed by an additional maximum mean depth cutoff of 67). Multi-allelic variants and indels were removed, resulting in 7,401 SNPs. Lastly, the 2,052 SNPs that were differentially fixed homozygote genotypes in the grandparents were used for creating the linkage map.

### Linkage map

A linkage map was constructed with JoinMap 4.0 (Van Ooijen 2006) using 212 F2 progeny derived from two F1 families. Out of the 224 genotyped individuals (including the 2 F0 and 4 F1), 2 F1 and 6 F2 were removed due to missing data (> 25%). Out of the 2,052 loci, homozygous for alternative alleles in the grandparents, we placed 1,597 in the final linkage map. Loci were excluded if positioned identical with another locus. Markers showing segregation distortion (χ^2^ test, P < 0.001) were excluded for linkage map reconstruction. Linkage groups were identified based on an independent logarithm of odds (LOD) threshold of 12. Unlinked markers were excluded. The strongest cross-link (SCL) in the final map is 5.4. The linkage map was built using the regression mapping algorithm, a recombination frequency smaller than 0.40, and a LOD larger than 3. Up to three rounds of marker positioning were conducted with a jump threshold of 5. A ripple was performed after the addition of each new marker. Map distances were calculated using the Kosambi mapping function. All markers resolved onto 22 linkage groups were matched to positions in the *Oreochromis niloticus* genome using a chain file (Brawand *et al*. 2014) with liftover (UCSC Genome Browser LiftOver tool; Hinrichs *et al*. 2006) to examine synteny of chromosomal locations and allow comparisons with other published studies.

### Anchoring of reference scaffolds

In order to reconstruct a chromosomal reference genome for *Pundamilia*, we used the linkage map to anchor the scaffolds of the *Pundamilia* genome from Brawand *et al*. (2014) onto the 22 *Pundamilia* linkage groups (Pun-LGs) identified during mapping (see paragraph above). We ordered and oriented the scaffolds with ALLMAPS (Tang *et al*. 2015). Gaps between the scaffolds were then estimated using interpolated recombination rate estimates based on the conversion between map distances (cM) and physical distances (bp) as implemented in the ALLMAPS function “estimategaps” (Tang *et al*. 2015). In addition to an improved reference version, resolving linkage groups, we compiled a chain file for converting positions on the original *Pundamilia nyererei* reference (Brawand *et al*. 2014) to our new reference (*Pundamilia* reference version 2.0) with ALLMAPS and in the opposite direction using chainSwap from kentUtils (https://github.com/ENCODE-DCC/kentUtils). We could then use the chain file to liftover the position of all 7,401 genotyped loci, using Picard liftoverVcf (http://broadinstitute.github.io/picard/index.html). In addition, we generated a new version of the NCBI *Pundamilia nyererei* RefSeq annotation file with the positions for reference version 2.0 by lifting over the positions from the NCBI PunNye1.0 annotation release 101 (https://www.ncbi.nlm.nih.gov/genome/annotation_euk/Pundamilia_nyererei/101/#BuildInfo) using the UCSC liftOver tool (Hinrichs *et al*. 2006) and custom-made chain files (see Table 2). By comparing physical (bp) and recombination distances (cM), we estimated recombination rates along the different linkage groups. First, we pruned the linkage map for markers generating negative recombination rates and markers that were less than 20 kb apart. Then we fitted a cubic smoothing spline to the physical (bp) and recombination (cM) distances using the R function “smooth.spline” setting the smoothing parameter (spar) to 0.7 and inferred the recombination positions in cM for the genomic positions as the first derivative of the “predict.smooth.spline” function.

### QTL mapping of sex

QTL mapping of the sex-determining region was performed with Rqtl (Broman *et al*. 2003) based on 209 individuals (3 F2 were discarded prior to analysis as they were juveniles) and 1,597 SNP markers. 137 males and 72 females were included. Sex was mapped by standard interval mapping as a binary trait and significance was determined by permutation (n = 1000). Bayesian confidence intervals were estimated as implemented in Rqtl and the highest LOD score was used to calculate the percent variance explained following 1 – 10^-2^ ^LOD/*n*^ (Broman and Sen 2009). Plotting phenotypic sex against the genotypes for the marker most strongly associated with sex, revealed two individuals labeled as females, but carrying a male genotype. Those individuals were dissected and their gonads were inspected, showing immature or undeveloped gonads indicating an error in phenotyping. The same plot also revealed both males (n = 74) and females (n= 32) that were heterozygous at the locus strongly associated with sex. To investigate if sex in those individuals was explained by another locus, we extracted the genotypes of these individuals and repeated the interval mapping. Further, we made use of 366 markers positioned on the linkage group containing the sex QTL and investigated segregation patterns at those loci in more detail in the larger of our mapping families (n = 122 F2 offspring). Based on the improved, annotated reference (v.2.0) we determined the number of annotated genes in the QTL interval and screened for candidate genes in sex determination.

### Data availability

All genomic resources (see Table 2) will be made available upon publication. Raw read sequencing files will be deposited on short read archive (fastq files for all 224 individuals). Genotype (vcf format) and phenotype file will also be made available.

## Results and Discussion

### Linkage map

The linkage map comprises 22 linkage groups containing 1,597 markers with an average marker distance of 1.01 cM adding up to a total map length of 1593.72 cM (Figure 1, Table 1). It is slightly longer than other maps published on Lake Victoria cichlids (1130.63 cM in Henning *et al*. 2014, 1133.2 cM in Kudo *et al*. 2015 and 1225.68 cM in Henning *et al*. 2017), but contains more markers with a lower average marker distance (1.30 cM (Henning *et al*. 2014) 1.09 cM (Henning *et al*. 2017) and 6.09 cM (Kudo *et al*. 2015)). The detection of 22 linkage groups is consistent with the expected number of chromosomes in haplo-tilapiine cichlids (Guyon *et al*. 2012). Out of 1,597 markers used to build the *Pundamilia* linkage map, 1,182 markers could be positioned onto *Oreochromis niloticus* linkage groups (Ore-LG). Figure 2 reveals extensive synteny between the chromosomes of these distantly related cichlid species. The linkage map presented here will facilitate comparative genomics and will enable comparisons of previous QTL results with newly established results (for an example see paragraph below on QTL for sex-determination) using Ore-LGs as a reference point.

**Table 1:** Summary of length and number of markers for each linkage group. Synteny between this study (Pun-LG) and the *Oreochromis niloticus* reference (Ore-LG) is indicated.

| *Pundamilia* LG | *Oreochromis* LG | length [cM] | # SNPs |
| --- | --- | --- | --- |
| Pun-LG1 | Ore-LG3 | 74.182 | 120 |
| Pun-LG2 | Ore-LG7 | 100.271 | 103 |
| Pun-LG3 | Ore-LG22 | 65.086 | 100 |
| Pun-LG4 | Ore-LG16-21 | 75.325 | 94 |
| Pun-LG5 | Ore-LG1 | 74.372 | 90 |
| Pun-LG6 | Ore-LG11 | 72.082 | 90 |
| Pun-LG7 | Ore-LG15 | 74.093 | 88 |
| Pun-LG8 | Ore-LG19 | 71.579 | 88 |
| Pun-LG9 | Ore-LG18 | 63.352 | 82 |
| Pun-LG10 | Ore-LG23 | 70.089 | 77 |
| Pun-LG11 | Ore-LG10 | 65.94 | 76 |
| Pun-LG12 | Ore-LG17 | 77.124 | 75 |
| Pun-LG13 | Ore-LG5 | 90.878 | 74 |
| Pun-LG14 | Ore-LG9 | 63.956 | 71 |
| Pun-LG15 | Ore-LG8-24 | 70.914 | 61 |
| Pun-LG16 | Ore-LG14 | 72.914 | 60 |
| Pun-LG17 | Ore-LG20 | 69.652 | 58 |
| Pun-LG18 | Ore-LG6 | 65.41 | 54 |
| Pun-LG19 | Ore-LG2 | 63.732 | 44 |
| Pun-LG20 | Ore-LG4 | 72.481 | 39 |
| Pun-LG21 | Ore-LG13 | 74.665 | 33 |
| Pun-LG22 | Ore-LG12 | 65.627 | 20 |
|  |  | 1593.724 | 1597 |

**Figure 1:**
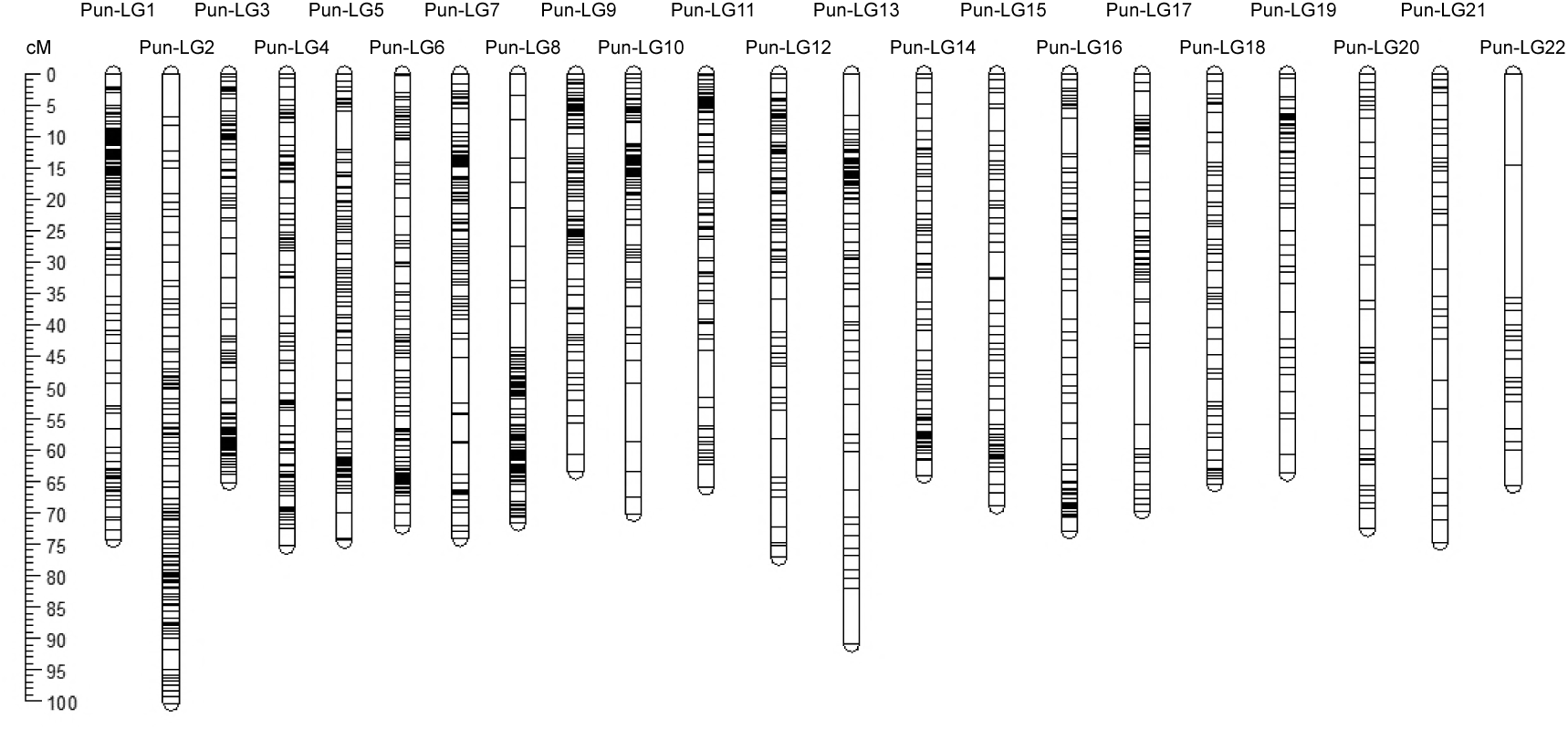
Linkage map indicating the positioning of 1,597 markers and Kosambi mapping length (cM) of 22 linkage groups.

**Figure 2:**
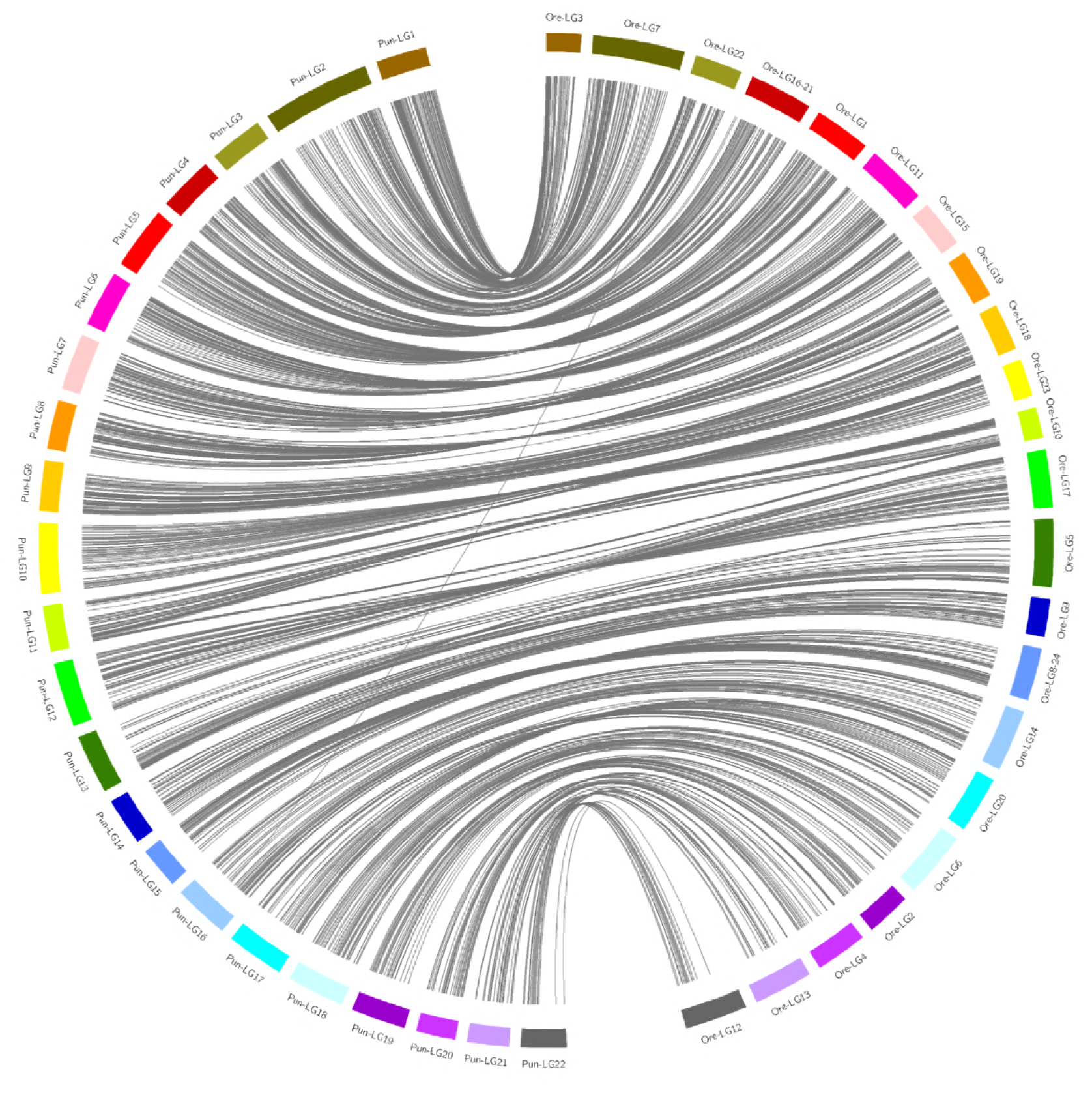
Synteny plot showing the correspondence of *Pundamilia* linkage groups (Pun-LG) with *Oreochromis niloticus* linkage groups (Ore-LG). Lines indicated markers used in linkage map construction, which could be positioned in the *Pundamilia* reference (v2.0) and lifted over to the *Oreochromis* reference.

### Improvement of the genomic resources for Lake Victoria cichlids (*Pundamilia*)

The *Pundamilia* linkage map provides a new chromosome framework for whole genome sequence assembly and map integration with more anchoring points then previous published maps. The anchored genome encompasses 78.7% of the total bases (653,642,680 bp) of the original *P. nyererei* reference genome based on 383 anchored scaffolds, of which 233 are now oriented. This is a slightly higher fraction than in the Lake Malawi cichlid *Metriaclima zebra*, where 564,259,264 bp (66.5%) of the genome sequence could be anchored to linkage groups (O’Quin *et al*. 2013). The mean marker density is 2.4 per megabase (Mb). The 6,853 remaining scaffolds could not be anchored due to lack of informative markers. This improved resolution of the new reference assembly (v2.0) will greatly facilitate genome scan approaches in Lake Victoria cichlids. Such approaches rely on the information from neighboring genomic positions to identify signatures of selection due to genetic hitchhiking. Any approaches evaluating or making use of linkage information, like linkage disequilibrium (LD) based genome scans, association studies or evaluations of the genomic landscape of divergence will now become feasible or more powerful. Together with the improved reference, we provide chain files to liftover positions from the previous version (v1.0) to the new chromosome level resolved reference version (v2.0). We further provide a matching annotation file based on the NCBI annotation (see Table 2 for a complete list of all genomic resources). Finally, we estimated recombination rates and show that those are highly variable across the genome ranging from 0 to 9.4 cM/Mb (Table 2), with a mean recombination rate of 2.3 cM/Mb. Knowledge of fine-scale patterns of recombination rate variation (see Figure 4C) will be useful for future studies of adaption and speciation (Stapley *et al*. 2017) in the exceptional species radiation of Lake Victoria cichlids.

**Table 2:**
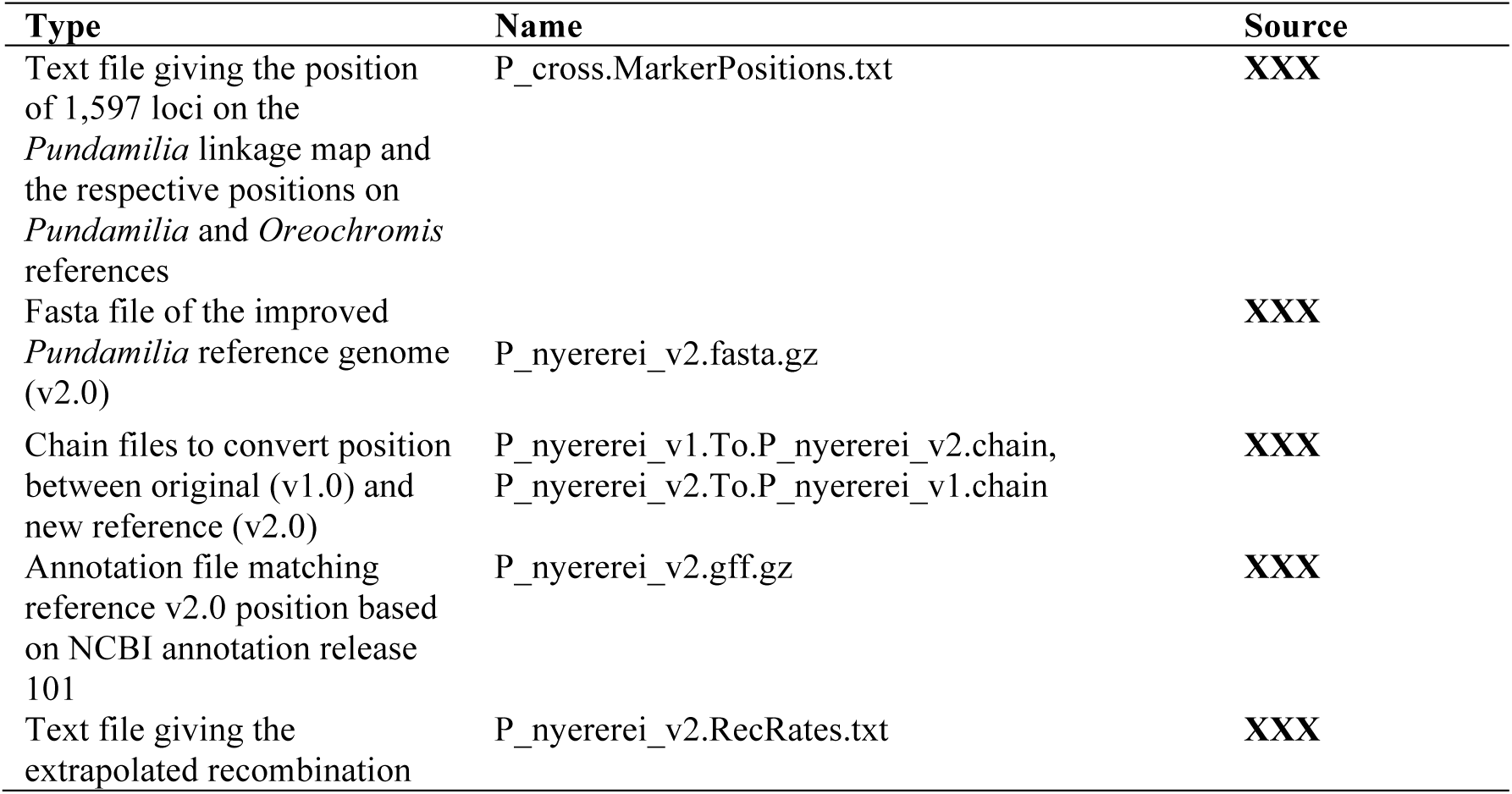
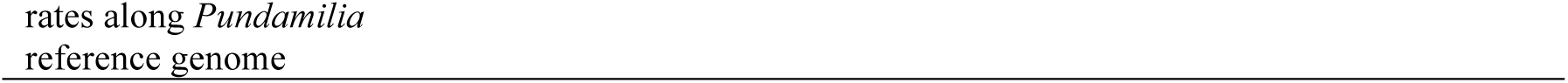
List of genomic resources provided with this manuscript

| Type | Name | Source |
| --- | --- | --- |
| Text file giving the position of 1,597 loci on the *Pundamilia* linkage map and the respective positions on *Pundamilia* and *Oreochromis* references | P_cross.MarkerPositions.txt | XXX |
| Fasta file of the improved *Pundamilia* reference genome (v2.0) | P_nyererei_v2.fasta.gz | XXX |
| Chain files to convert position between original (v1.0) and new reference (v2.0) | P_nyererei_v1.To.P_nyererei_v2.chain,<br>P_nyererei_v2.To.P_nyererei_v1.chain | XXX |
| Annotation file matching reference v2.0 position based on NCBI annotation release 101 | P_nyererei_v2.gff.gz | XXX |
| Text file giving the extrapolated recombination | P_nyererei_v2.RecRates.txt | XXX |

### Characterization of sex-determination in *Pundamilia*

Our knowledge of sex determination in *Pundamilia*, a prime model system of sympatric speciation in Lake Victoria, had been limited. Here, we mapped sex to Pun-LG10, which is homologous with Ore-LG23 (Figure 3; p << 0.001, LOD = 26.5). We did not find any further associations on any of the other LGs. Ore-LG23 has been previously identified as one potential sex-determining LG in *Oreochromis* (Cnaani *et al*. 2008; Palaiokostas *et al*. 2013) and in four cichlid tribes from Lake Tanganyika overexpression of male specific genes accumulate on Ore-LG23 (Böhne et al. 2014). Early work on sex determination in Lake Victoria cichlids had suggested polymorphisms at several unlinked genomic regions to be associated with sex, and invoked a major effect locus and some modifiers (Seehausen et al. 1999). Recent QTL mapping identified genomic regions involved in sex determination in Lake Victoria cichlids on Ore-LG5 and Ore-LG2 (Kudo *et al*. 2015) or on derived, female specific B chromosomes (Yoshida *et al*. 2011). Ore-LG5 was repeatedly found to be involved in sex-determination in other cichlids, e.g. in the riverine haplochromine cichlids *Astatotilapia burtoni* and *Astatotilapia calliptera* from Lakes Tanganyika and Malawi and associated rivers (Roberts *et al*. 2016; Böhne *et al*. 2016; Peterson *et al*. 2017), in *Cyprichromis leptosoma* from Lake Tanganyika (Gammerdinger *et al*. 2018) and in *Labeotropheus trewavasae* and across some *Metriaclima* species from Lake Malawi (Ser *et al*. 2010; Parnell and Streelman 2013).

**Figure 3:**
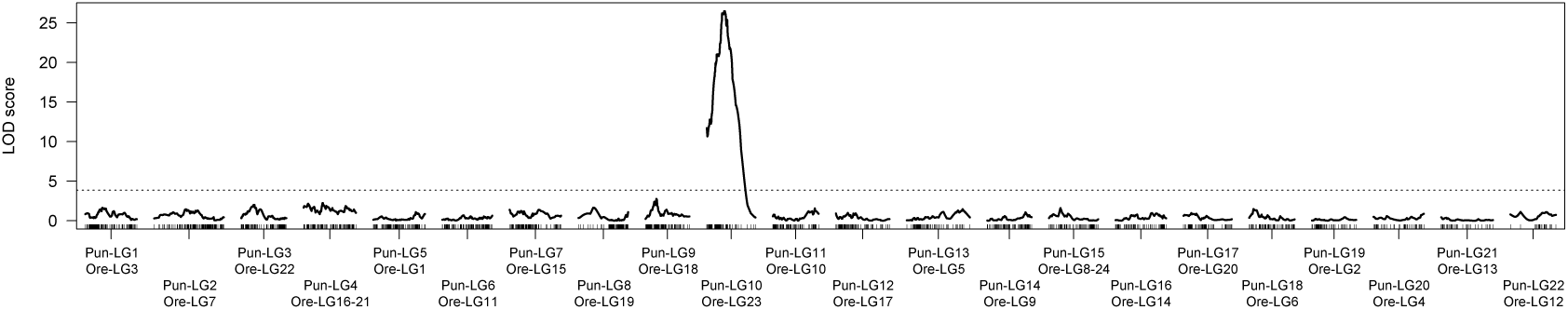
QTL mapping of sex. LOD scores across the 22 linkage groups are shown. Genome-wide significance levels are indicated by horizontal lines (alpha = 0.05 dotted line). Marker loci are indicated along the x-axis.

The mapping interval (Bayesian confidence interval of 5.7 cM, 21.7 to 27.4 cM; Figure 4B) in total covers four markers and spans a region of ∼1.9 Mb (Figure 4C). The marker showing the strongest association with sex in our study (Figure 4A) explains 44% of the phenotypic variance in sex. Sex is not entirely explained by this marker as we had misidentified two likely sub-adults (gonads appear not developed at hindsight inspection) as females, and due to 106 individuals, both males and females, which are heterozygous at this position (Figure 4A). Repeating the mapping procedure for those individuals again identified a region on Pun-LG10 (Ore-LG23) as weakly associated with sex (p = 0.177, LOD = 3.33, position right to previous interval at 28.8 cM). This suggest that none of the markers used to build the linkage map is determining sex directly, but that the causal locus can be found close by and indicates that there are no further major genetic determines of sex segregating in this cross. Investigating the segregation patterns in the larger of the F2 mapping-families (n = 122) more in detail, the loci selected to built the map, reciprocal homozygous in F0 female (AA) and male (BB) and heterozygous in both F1 (AB), segregate as expected in a 50:50 ratio of AA:AB in F2 females and AB:BB in F2 males (Figure 5A). However, evaluating segregation patterns of the additional markers genotyped but not used for the construction of the linkage map, indicate that the sex determination system on Pun-LG10 is male heterogametic (XY, Figure 5B). We identified 57 loci between 0 and 35 Mb that were homozygous in the F0 and F1 females and heterozygous in the F0 and F1 males; these markers are similarly homozygous in all F2 females and heterozygous in all F2 males, consistent with females being XX and males being XY (Figure 5B, the plot also shows 13 loci > 35 Mb). Additional evidence comes from markers heterozygous in the F0 female (AB) and homozygous in the F0 male (BB), for which we find all 35 loci for positions < 33 Mb heterozygous (AB) for both F1 individuals. The heterozygous loci in both F1 are a segregation pattern only consistent with male heterogametic (XY) segregation. If females would be heterogametic (ZW) those loci would need to be homozygous (BB) in one of the F1 and not heterozygous (AB) in both as observed in that 33 Mb region. Sex-averaged recombination rates around the QTL are low and even close to zero within 20 Mb proximity to the mapping interval (Figure 4C). Such a pattern, potentially due to suppressed recombination in the heterogametic sex (males), might indicate initial steps toward the evolution of a heteromorphic (degenerated) sex (Y) chromosome (Charlesworth, 1991).

**Figure 4:**
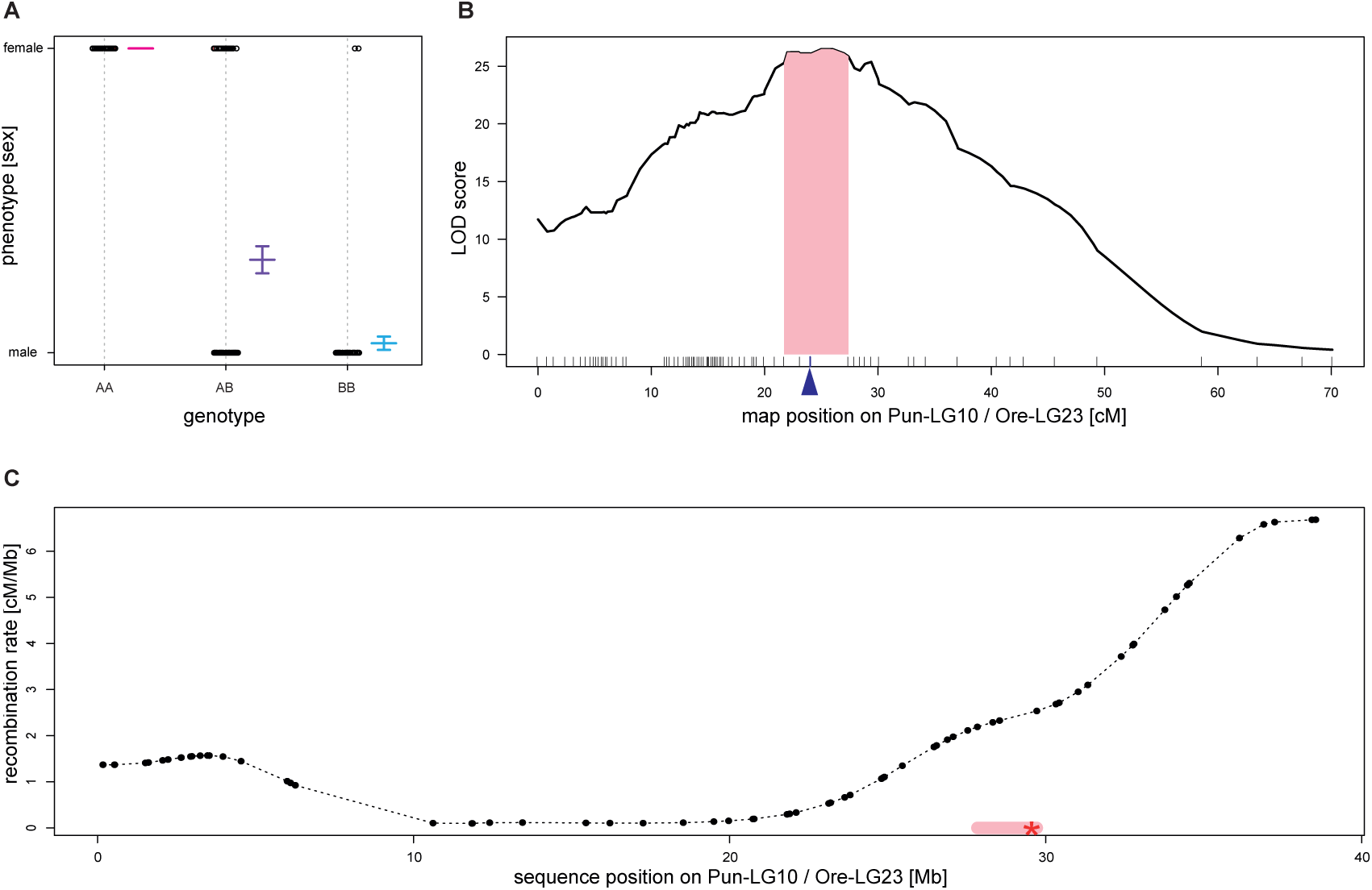
A) Phenotypic effect of genotypes at the locus most strongly associated with sex. The plot identifies two females as likely phenotypic errors and 106 individuals heterozygote at that locus. B) Plot of LOD scores indicating the region of strongest association with sex on Pun-LG10 (Ore-LG23). The Bayesian confidence interval is highlighted in light pink. Marker loci are indicated along the x-axis. The locus shown in panel A is indicated by a blue arrow. C) Variation in recombination rates (sex-averaged) along Pun-LG10 (Ore-LG23). The Bayesian confidence interval (pink highlight) is situated next to a region of low recombination. The red star indicates the position of *amh* (candidate gene for sex determination).

**Figure 5:**
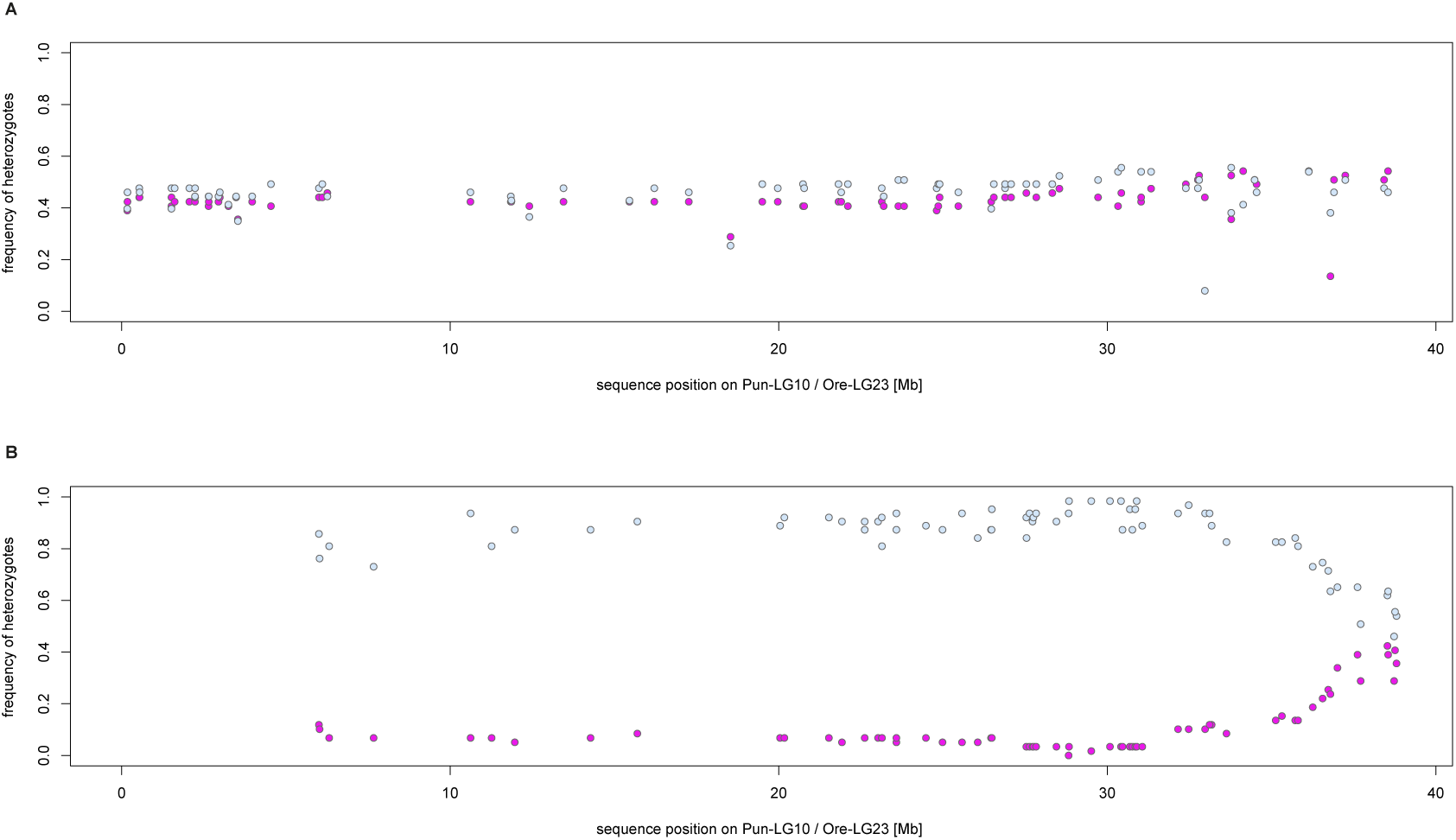
Frequency of heterozygote individuals (n = 122), separated by sex (63 males: light blue, 59 females: light pink) for markers selected by their segregation pattern in the larger mapping family and their position on Pun-LG10 (Ore-LG23). A) 78 markers, selected as reciprocally homozygous (AA/BB) in the F0 and heterozygote in both F1 (AB/AB), segregate as expected in a 50:50 ratio of AA:AB in F2 females and AB:BB in F2 males, resulting in frequency of heterozygous F2 individuals around 0.5 for both sexes. B) 70 markers, selected as homozygote in the F0 and F1 females and heterozygote in the F0 and F1 males, segregate similarly in the F2; i.e. the frequency of heterozygous individuals is approaching 0 in females and 1 in males for positions < 35 Mb.

Within our mapping interval of ∼1.9 Mb, 65 genes, based on the NCBI annotation for the new *Pundamilia* reference assembly, can be found (Table S1). Among them is the anti-müllerian hormone (*amh)*, a master gene for sex determination in other fish. *Amh* is part of the transforming growth factor beta pathway, responsible for the regression of Müllerian ducts in tetrapods (Josso *et al*. 2001). Even though teleost fish do not have Müllerian ducts, the *amh* pathway has a prominent role in sex determination for several distantly related fish species. In the Japanese pufferfish (*Takifugu rubripes*), a mutation in the receptor of the *amh* (amhrII) determines sex (Kamiya *et al*. 2012). The *amhy* (Y chromosome-specific anti-müllerian hormone) gene has been inserted upstream of *amh* in the cascade of male development in the neotropical silverside *Odonthestes hatcheri* (Hattori *et al*. 2012). Similarly, in *Oreochromis niloticus*, a Y-linked duplicate of *amh* acts as a major sex determination locus (Eshel *et al*. 2012; Li *et al*. 2015). In *Oryzias luzonensis*, a mutation of an *amh* related ligand gsdf^y^ is responsible for sex determination (Myosho *et al*. 2012). The same ligand is suggested to be involved in sex determination in the haplochromine cichlid *Astatotilapia calliptera* (Peterson *et al*. 2017). Beside the two master sex determination genes in *Oreochromis niloticus* on LG23 (*amh*) and in *Astatotilapia calliptera* on LG7 (*gsdf*) (Peterson *et al*. 2017), candidates for sex determination in cichlids have not been shown to be directly involved in sex determination in other species (Heule *et al.* 2014b, Böhne *et al.* 2016, Gammerdinger *et al.* 2018, but see Böhne *et al*. 2014). They might in this matter act as so-called “newcomers” (Herpin and Schartl 2015). Our results indicate that in the Lake Victoria cichlid *Pundamilia* Pun-LG10 (Ore-LG 23) acts as an (evolving) sex chromosome, even though it might not be the only region controlling sex in *Pundamilia*. The anti-müllerian hormone *amh* (or a derived copy) appears to be a very good candidate influencing sexual development in *Pundamilia*, but further work is warranted to characterize the genomic candidate region and the impact of this candidate gene on sex determination.

A recent meta-analysis showed that transitions between sex determination systems are frequent across various fish species, including transitions to and between heteromorphic sex chromosomes (Pennell *et al.* 2018). In cichlids a high turnover of sex determination systems was described in Lake Malawi (Ser *et al*. 2010), Lake Tanganyika (Böhne *et al*. 2014; Gammerdinger *et al*. 2018), and oreochromine cichlids (Cnaani *et al*. 2008). The circumstance that *amh*, Pun-LG10 or a homologous region was not invoked in sex determination in other Lake Victoria cichlids that have previously been used for mapping sex (Kudo *et al*. 2015; Yoshida *et al.* 2011) implies that multiple sex determining systems segregate among the species of Lake Victoria cichlid fish as well. This is consistent with early work on sex determination in this group (Seehausen *et al*. 1999). Given the extreme youth of the Lake Victoria species radiation (∼15,000 years; Seehausen 2006), this may be surprising at first. Recent work, however, has shown that much of the genetic variation in the radiation is much older than the species radiation and took its origin in a hybridization event between two anciently divergent cichlid lineages from which all 500+ species of the radiation evolved (Meier *et al*. 2017a). It is tempting to speculate that the variation in sex determination systems between and within species of this radiation traces its roots to these ancient lineages too, something that can now be tested.

---

Table S1: List of genes overlapping the mapping interval on Pun-LG10 (Ore-LG23). Gene position as annotated in *Pundamilia* (v2.0) annotation file based on NCBI annotations using Uni-Prot.

## Acknowledgements

We thank Katie Peichel for her critical input during the analysis of the data and manuscript preparation. We thank Andreas Taverna, Alan Smith, and Ola Svensson for fish maintenance and breeding. We thank David Marques for lifting over the annotation file. We thank the team from the Genetic Diversity Center (GDC) at ETH Zürich for bioinformatics support. Genomic analyses were performed using the computing infrastructure of the GDC and the Euler computer cluster at ETH Zurich. JS was supported by the German Research Foundation grant SCHW-1690/1-1.

## Author’s contributions

PGDF, JS, MPH, and JIM performed the experiment and the analysis. OS conceived the original idea and supervised the project. PGDF and JS took the lead in writing the manuscript. All authors provided critical feedback and helped shape the research, analysis, and manuscript.

